# Structural basis of substrate recognition and catalysis by fucosyltransferase 8

**DOI:** 10.1101/2020.02.14.949818

**Authors:** Michael A. Järvå, Marija Dramicanin, James P. Lingford, Runyu Mao, Alan John, Kate Jarman, Rhys W. Grinter, Ethan D. Goddard-Borger

**Affiliations:** The Walter and Eliza Hall Institute of Medical Research, Parkville, Victoria 3052, Australia; Department of Medical Biology, University of Melbourne, Parkville, Victoria 3010, Australia; Department of Microbiology, Monash Biomedicine Discovery Institute, Monash University, Clayton, VIC 3800, Australia

**Keywords:** glycosyltransferase, N-glycosylation, core fucose, enzyme mechanism, structural biology

## Abstract

Fucosylation of the inner-most N-acetyl-glucosamine (GlcNAc) of N-glycans by fucosyltransferase 8 (FUT8) is an important step in the maturation of complex and hybrid N-glycans. This simple modification can have a dramatic impact on the activity and half-life of glycoproteins. These effects are relevant to understanding the invasiveness of some cancers, the development of monoclonal antibody therapeutics, and to a congenital disorder of glycosylation. The acceptor substrate preferences of FUT8 are well characterised and provide a framework for understanding N-glycan maturation in the Golgi, however the structural basis for these substrate preferences and the mechanism through which catalysis is achieved remains unknown. Here, we describe several structures of mouse and human FUT8 in the apo state and in complex with guanosine diphosphate (GDP), a mimic of the donor substrate, and a glycopeptide acceptor substrate. These structures provide insights into: a unique conformational change associated with donor substrate binding; common strategies employed by fucosyltransferases to coordinate GDP; features that define acceptor substrate preferences; and a likely mechanism for enzyme catalysis. Together with molecular dynamics simulations, the structures also reveal how FUT8 dimerisation plays an important role in defining the acceptor substrate binding site. Collectively, this information significantly builds on our understanding of the core-fucosylation process.

## Introduction

Fucosyltransferase 8 (FUT8) is the mammalian α-1,6-fucosyltransferase responsible for modifying the inner-most (reducing-end) GlcNAc of hybrid and complex N-glycans. This modification, referred to as core-fucosylation, is ubiquitous throughout mammalian tissues and represents an important step in the maturation of complex N-glycans within the Golgi apparatus. Core fucosylation modulates the activity of many cell surface receptors, including: TGFβ1R^1, 2^, EGFR^3^, BCR^4^, TCR^5, 6^, CD14-mediated TLR2/4 signalling^7, 8^, and PD-1^9^. It also modulates the affinity of ligands for their receptors, the most notable example being the role that core fucose plays in decreasing the affinity of immunoglobulin G (IgG) for FcγRIIIa^10, 11^. This latter phenomenon has inspired the development of next-generation therapeutic monoclonal antibodies that more effectively engage FcγRIIIa and demonstrate superior antibody-dependent cellular cytotoxicity (ADCC)^11, 12^. REcently, dectin-I was identified as the first endogenous lectin that specifically recognises core fucose^13^. Colelctively, these and other findings demonstrate that FUT8 as plays a central role in modulating the activity of many cell-surface receptors.

Within mice, loss of function mutations in FUT8 result in severe growth retardation and the development of an emphesema-like lung phenotype, purportedly due to dysregulation of TGF-β1 and EGFR signalling^1, 14^. These animals also exhibited behavioural abnormalities^15^. Many of these phenotypes are also observed in patients with the recently described FUT8 congenital disorder of glycosylation (CDG-FUT8)^16^. In contrast to CDG-FUT8, which features the ablation of FUT8 activity, many cancers upregulate FUT8 expression and this correlates with a poor prognosis^17^. In melanomas, increased FUT8 activity stabilises L1CAM to promote metastasis^18^. Metastasis is also promoted by FUT8 in breast cancers, where increased core-fucosylation of TGFβ1R promotes strong constitutive signalling through this receptor and tumour cell migration^19^. The increased core fucosylation of α-fetoprotein is also a well-established biomarker of hepatocellular carcinoma (HCC)^20^.

Some have speculated that FUT8 antagonists may have therapeutic potential for the treatment of cancer^9,18^, though questions remain around how a hypothetic FUT8 antagonist might impact host immune responses to tumour cells. Regardless, no drug-like small molecule inhibitors have yet been reported for FUT8, or any other human fucosyltransferase (FUT). To some degree, drug discovery efforts are impeded by a limited structural understanding of this enzyme and the mechanism it employs to perform core fucosylation. The only reported FUT8 structure possesses no bound ligands,^21^ and our only insights into donor and acceptor substrate binding come from STD-NMR, molecular dynamics and docking studies^22, 23^. To gain a thorough understanding of how FUT8 recognises both its donor and acceptor substrates to catalyse core fucosylation, we revisited the structural biology of FUT8. The structures we obtained provide fresh insights into the conformational dynamics and molecular interactions associated with catalysis.

## Results

### Structural insights into nucleotide recognition by FUT8

A truncated human FUT8 (HsFUT8_105-575_) construct, which is missing the N-terminal transmembrane domain and unstructured region, was expressed in Sf21 insect cells. The activity of the purified protein was verified using the GDP-Glo™ glycosyltransferase assay with an asialo-agalacto-biantennary glycopeptide (A2SGP) derived from chicken eggs as an acceptor substrate (**Figure 1A**). Using this assay, we determined a K_M_ of 4.2 μM for GDP-fucose (GDP-Fuc) and 12 μM for A2SGP (**Figure 1B**), which was in broad agreement with previously reported values^24^.

**Figure 1.**
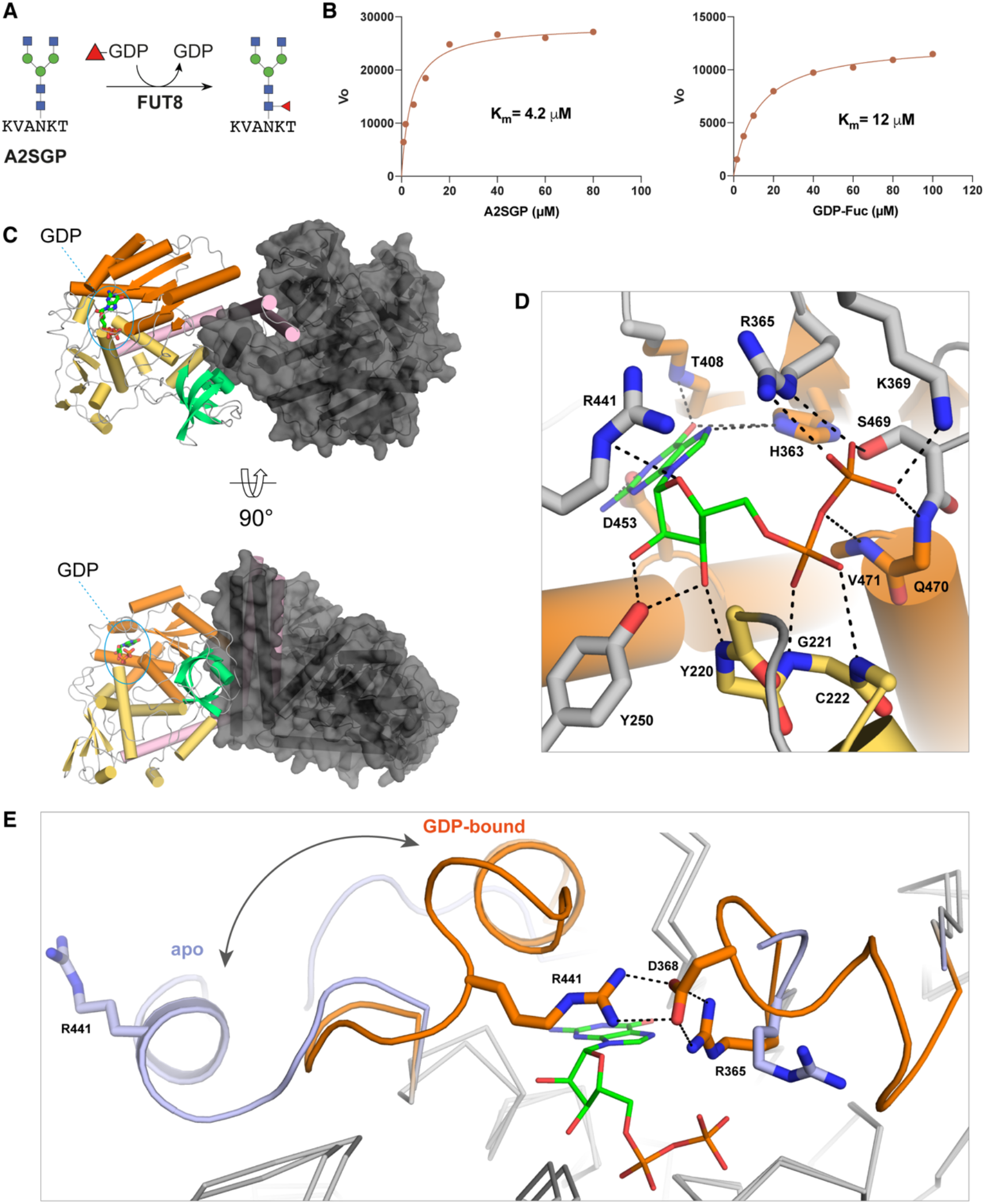
Structures of mouse and human FUT8 with and without GDP bound. (A) HsFUT8 is active and has K_M_ values that are in agreement with those previously reported^24^. (B) The domain structure of FUT8 (coiled coil = pink, Rossman = orange and yellow, SH3 = teal) and the interactions between each the two molecules in the asymmetric unit. (C) The hydrogen-bonding interactions between MmFUT8 and GDP (see also Figure S4). (D) An overlay of the GDP-binding sites of apo-HsFUT8 (blue) and GDP-bound MmFUT8 (orange), illustrating the conformational changes observed for loops A and B. A salt bridge between D368 and R365/441 from loops A and B, respectively, forms upon encapsulation of GDP.

Since glycosyltransferases rapidly hydrolyse their sugar nucleotide donor substrates on a protein crystallisation time-scale, we attempted to co-crystallise HsFUT8 with GDP rather than GDP-Fuc. These attempts failed to provide any crystals of the complex, though we did obtain crystals of the apo form that enabled a re-determination of the unliganded structure at higher resolution (2.28Å) than the existing structure (2.61 Å for PDB ID 2DE0) with superior refinement statistics (**Table 1**)^21^. As an alternative approach, we cloned and expressed the mouse FUT8 (MmFUT8_68-575_) using a similar method as for the human protein. MmFUT8 is 96.6% identical to the human homologue over the length of this truncated construct (**Figure S1**). Extensive crystallisation screens with this slightly different protein and GDP provided crystals that yielded a structure of MmFUT8 in its apo (1.80Å) and GDP-bound (2.50Å) forms (**Table 1**).

**Table 1.** Refinement statistics for the structures reported in this study.

| Structure | apo-MmFUT8 | MmFUT8–GDP | apo-HsFUT8 | HsFUT8–GDP–<br>A2SGP |
| --- | --- | --- | --- | --- |
| PDB ID: | 6VLF | 6VLG | 6VLE | 6VLD |
| Space group | C 1 2 1 | P 6<sub>5</sub> 2 2 | P 1 2<sub>1</sub> 1 | C 1 2 1 |
| No of protein chains in AU | 2 | 4 | 2 | 4 |
| Cell dimensions |  |  |  |  |
| *a*, *b*, *c* (Å) | 184.87, 71.34, 126.95 | 150.82, 150.82, 472.14 | 95.02, 62.20, 109.90 | 208.31, 68.45, 249.98 |
| *a*, *b*, *g* (°) | 90, 126.08, 90 | 90, 90, 120 | 90, 90.88, 90 | 90, 111.21, 90 |
| Wavelength (Å) | 0.9537 | 0.9537 | 0.9537 | 0.9537 |
| Resolution (Å)* | 49.20-1.80 (1.83-1.80) | 49.37-2.50 (2.54-2.50) | 47.50-2.28 (2.34-2.28) | 46.65-2.28 (2.32-2.28) |
| *R*<sub>sym</sub> or *R*<sub>merge</sub>* | 0.047 (1.167) | 0.150 (0.664) | 0.120 (1.520) | 0.095 (1.347) |
| *R*<sub>pim</sub>* | 0.044 (1.055) | 0.066 (0.366) | 0.092 (1.143) | 0.066 (0.949) |
| *I* / *sI* * | 10.6 (0.9) | 9.5 (2.0) | 8.2 (1.0) | 10.2 (1.1) |
| CC(1/2) | 0.999 (0.463) | 0.995 (0.811) | 0.997 (0.630) | 0.998 (0.435) |
| Completeness (%)* | 99.6 (99.7) | 98.8 (85.8) | 99.8 (100) | 99.9 (99.9) |
| Redundancy* | 3.4 (3.5) | 10.4 (6.2) | 5.0 (5.1) | 5.7 (5.6) |
| Wilson B-factor (Å<sup>2</sup>) | 31.78 | 33.56 | 40.13 | 47.03 |
| **Refinement** |  |  |  |  |
| Resolution (Å) | 46.42-1.80 | 49.37-2.50 | 47.26-2.28 | 48.66-2.28 |
| No. reflections | 123,188 | 108,979 | 58,849 | 150,217 |
| *R*<sub>work</sub> / *R*<sub>free</sub> | 0.1842 / 0.2082 | 0.1822 / 0.2273 | 0.2019 / 0.2340 | 0.1914 / 0.2239 |
| No. non-hydrogen atoms |  |  |  |  |
| Protein | 7384 | 15248 | 7492 | 15083 |
| GDP | n/a | 112 | n/a | 56 |
| A2SGP | n/a | n/a | n/a | 392 |
| Water | 626 | 925 | 376 | 628 |
| *B*-factors |  |  |  |  |
| Protein | 43.4 | 43.3 | 50.9 | 64.2 |
| GDP | n/a | 32.0 | n/a | 56.2 |
| A2SGP | n/a | n/a | n/a | 65.6 |
| Water | 47.9 | 43.0 | 48.5 | 55.3 |
| R.m.s. deviations |  |  |  |  |
| Bond lengths (Å) | 0.005 | 0.003 | 0.001 | 0.002 |
| Bond angle (°) | 0.72 | 0.61 | 0.417 | 0.482 |
| Ramachandran plot (%) |  |  |  |  |
| Favored | 97.62 | 96.73 | 97.8 | 96.58 |
| Allowed | 2.38 | 3.22 | 2.2 | 3.42 |
| Disallowed | 0 | 0 | 0 | 0.05 |

The overall fold of these three FUT8 crystal structures is as previously described^21^: an N-terminal coiled-coil domain, followed by two Rossman folds forming a GDP-substrate-binding site, and a C-terminal SH3-domain of unknown function, as illustrated for MmFUT8-GDP in **Figure 1C**. The backbone RMSD between the four structures is low, <0.4Å, (**Figure S2**). The most notable backbone perturbations observed are for two loops, Arg365-Ala375 (loop A) and Asp429-Asn446 (loop B), which are disordered or displaced in the apo-MmFUT8 and apo-HsFUT8 structure but become ordered and completely encapsulate GDP upon binding (**Figure 1D,E**). This reorganisation involves the creation of several new interactions between both loops, most notably a salt bridge between Asp368 and Arg365 of loop A and Arg441 of loop B (**Figure 1E**). Arg365 also forms a salt bridge with the beta-phosphate of GDP, providing a link between ligand binding and organisation of the encapsulating loops (**Figure 1D,E**). Mutation of Arg365 to Ala abolishes FUT8 activity^25^, confirming that this residue plays a key role in organising the encapsulating loops around the nucleotide. A detailed list of all GDP-FUT8 hydrogen bonds is provided in **Figure S3**. Other noteworthy interactions include those between Asp453/His363 and the guanine base and the interactions between the ribose hydroxyl groups and Tyr250 (**Figure 1D**).

The conformational change associated with GDP encapsulation by FUT8 is unique amongst the FUTs studied to date. It results in burial of 96% of GDP’s surface area (**Figure S3**). This is comparable to, or slightly higher than, that observed for other FUT:GDP complexes, including: AtFUT1 (95%)^26^, NodZ (83%)^27^, CePOFUT1 (90%)^28^, MmPOFUT1 (86%)^29^ and CePOFUT2 (95%)^30^ (**Figure 2**). Remarkably, the conformational pose of the GDP ligand and GDP-FUT interactions are nearly identical in all FUTs, including FUT8, despite significant divergence in sequence and domain architecture. Residues analogous to FUT8’s Ser469, Arg365, Asp453, and His363 are conserved at a structural level across all FUTs (**Figure 2**). This observation will have important consequences for the development of competitive inhibitors of the FUTs.

**Figure 2.**
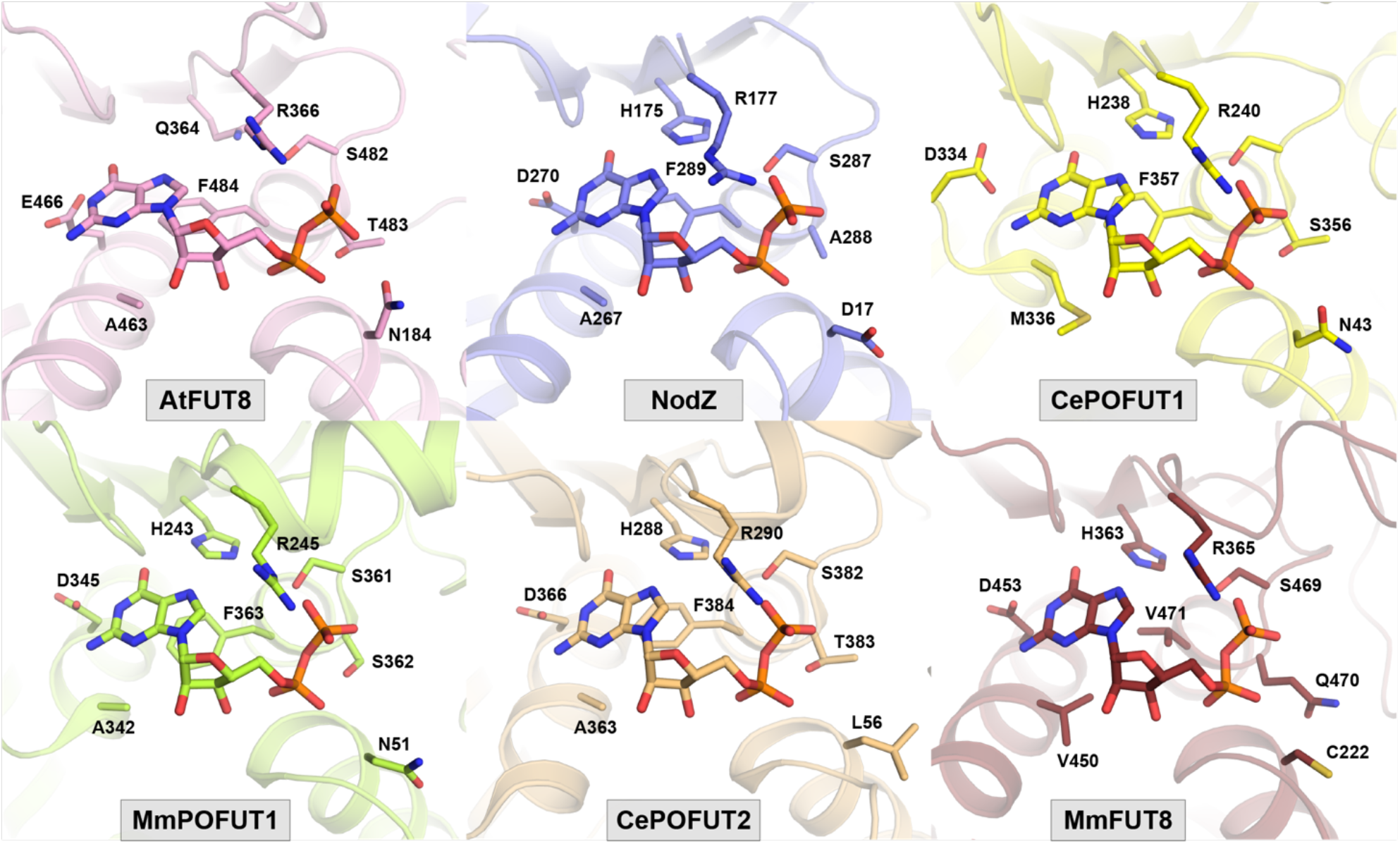
Conserved residues defining the GDP-binding site across all known structures of fucosyltransferases in complex with GDP (PDB ID for AtFUT1: 5KWK^26^; NodZ: 3SIW^27^; CePOFUT1: 3ZY3^28^; MmPOFUT1: 5KXQ^29^; CePOFUT2: 5FOE^30^; MmFUT8: 6VLG).

### Structural insights into N-glycan acceptor substrate recognition by FUT8

Co-crystallisation of HsFUT8_105-575_ with GDP and the A2SGP glycopeptide acceptor substrate provided crystals that yielded a structure with four HsFUT8 monomers in the asymmetric unit. All molecules were bound to A2SGP but only one molecule in each dimer pair also bound GDP. As such, this structure provided information for the FUT8:GDP:A2SGP ternary complex and the FUT8:A2SGP binary complex. The apparent ability of GDP and A2SGP to bind FUT8 independently of each other is consistent with a rapid equilibrium random mechanism, which has been previously been established for FUT8^24^.

All sugars of the A2SGP substrate, and the Asn side chain to which they were attached, were resolved and modelled for each monomer (**Figure 3A**). Upon binding to FUT8, the N-glycan buries 44% of its surface area. Hydrogen-bonding interactions between FUT8 and A2SGP are almost exclusively between the enzyme and the GlcNAc units comprising the core chitobiose unit and non-reducing ends of the bisected glycan (**Figure 3A,B**): the mannose units do not make any notable hydrogen-bonding interactions with FUT8. For the two protein chains in the assymetric unit without GDP bound, loops A and B remain disordered, as for the apo structures. However, in the ternary complex with GDP bound, these loops encapsulate GDP, as observed for the MmFUT8-GDP structure. Inspection of the region between the beta-phosphate of GDP and the 6-hydroxyl group of the innermost GlcNAc residue of A2SGP provides insights into which residues play a role in catalysis. Glu373 forms an intimate hydrogen bond (2.3 Å) with the 6-hydroxyl group and also interacts with Lys369, which in turn forms an intimate contact with the beta-phosphate of GDP (**Figure 3C**). This suggests that Glu273 acts as the catalytic base for catalysis and is capable of relaying a proton through Lys369 to the departing beta-phosphate of GDP. In this way, the Glu273/Lys369 pair act as proton conduit and catalytic base/acid, respectively.

**Figure 3.**
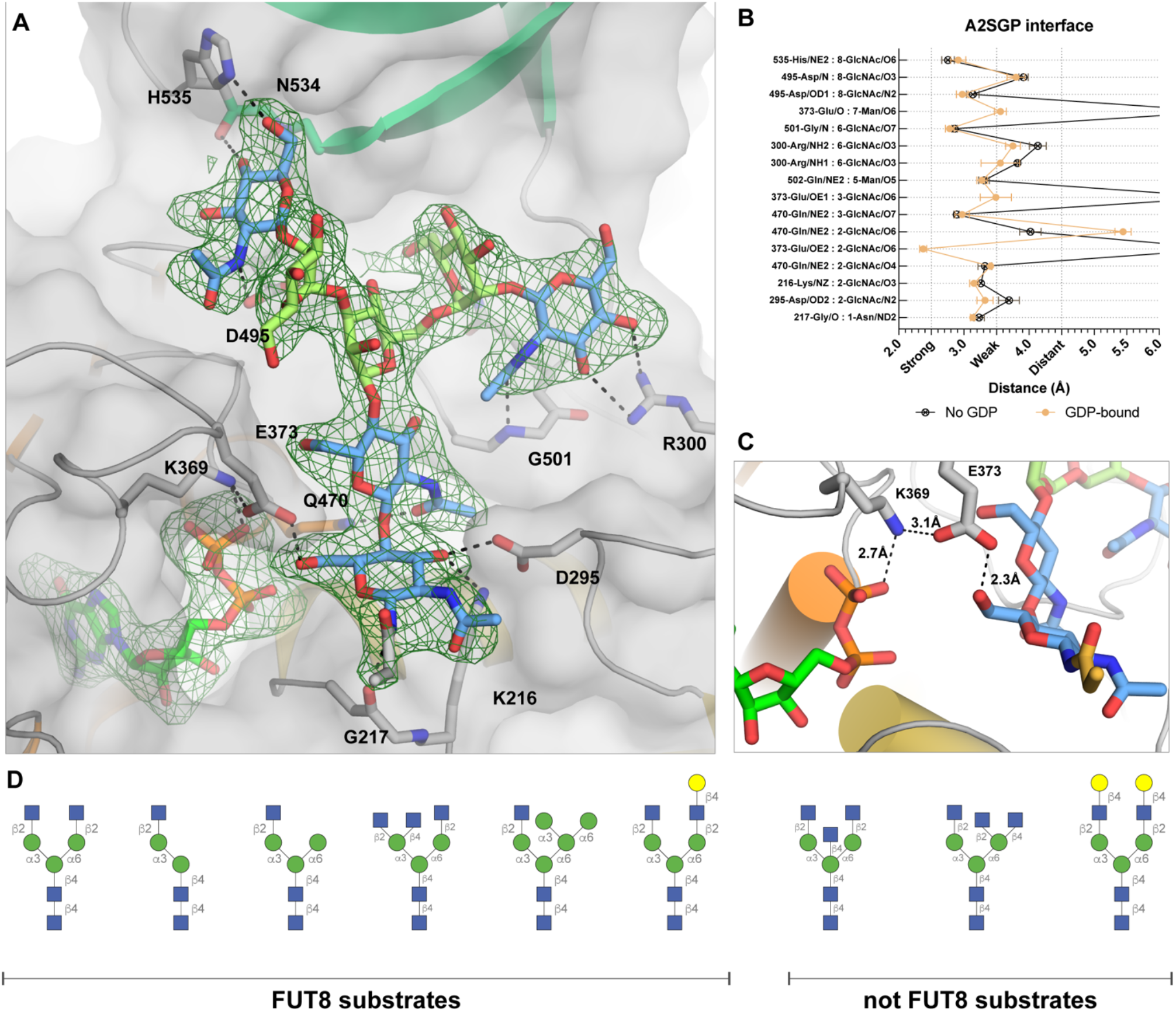
Acceptor substrate recognition by FUT8. (A) Interactions between FUT8 and the A2SGP N-glycan acceptor substrate, with an Fo-Fc omit map contoured at 1.5σ around the N-glycan and GDP. Residues making hydrogen bond interaction to the N-glycan are indicated. SH3 domain is coloured in teal. (B) A list of the interactions and hydrogen bond distances between A2SGP and FUT8. (C) A close up of the active site illustrating a potential role for E373 and K369 as a proton relay to facilitate electrophilic migration of fucose from GDP-Fuc to the 6-hydroxyl group of the innermost GlcNAc. (D) A selection of N-glycans known to be modified or not modified by FUT8^31, 32^, for comparison to the structure depicted in (A).

Two previous studies have explored the acceptor substrate specificity of FUT8 to ascertain what features promote or impair FUT8 activity^31, 32^ and these results are summarised in **Figure 3D**. It is clear through comparison to our structure that a bisecting GlcNAc would be sterically occluded by the SH3 domain of FUT8 (**Figure 3A**), consistent with this modification’s ability to block core-fucosylation^31, 32^. Modifications of the α6-branch of the glycan are well-tolerated by FUT8, and it is clear from our structure that there is sufficient space to accommodate most types of truncation, elongation or branching at this position. The one exception is elongation with a terminal β-1,4-GlcNAc, which would introduce steric clashes with the SH3 domain. Modification of the α3-branch is not well-tolerated by FUT8, and all typical elongation or branching residues (**Figure 3D**) introduce steric clashes that would preclude binding. Notably, FUT8 activity requires the terminal β-1,2-GlcNAc of the α3-branch, suggesting that the intimate hydrogen bond between His353 and the 6-hydroxyl group of this GlcNAc is an important contributor to acceptor substrate binding.

### FUT8 dimerisation and orientation of SH3-domain for acceptor substrate recognition

In the asymmetric units of all structures determined here, MmFUT8 and HsFUT8 formed an apparent dimer through the formation of a four-helix bundle from their N-terminal coiled-coil domains. These helices interact with their neighbour’s SH3-domain (**Figure 4A**). This dimer could be observed in the previously determined apo structure of FUT8^21^, yet the structure was reported on as a monomer. Other publications have reported that HsFUT8 is a monomer in solution, based on size exclusion chromatography experiments^33^, and this has influenced the way in which molecular dynamic and docking simulations of FUT8 have been conducted^22, 23^, potentially compromising their conclusions. To address this inconsistency in the literature, we performed SEC-SAXS on FUT8 (**Figure 4B,C**), which conclusively demonstrated that FUT8 exists as a dimer in solution. The buried surface area for dimerisation was similar for all four structures reported here (**Table S4**), irrespective of what ligands were bound, and involves the same residues forming inter-chain salt bridges and hydrogen bonds (**Figure 4A**).

**Figure 4.**
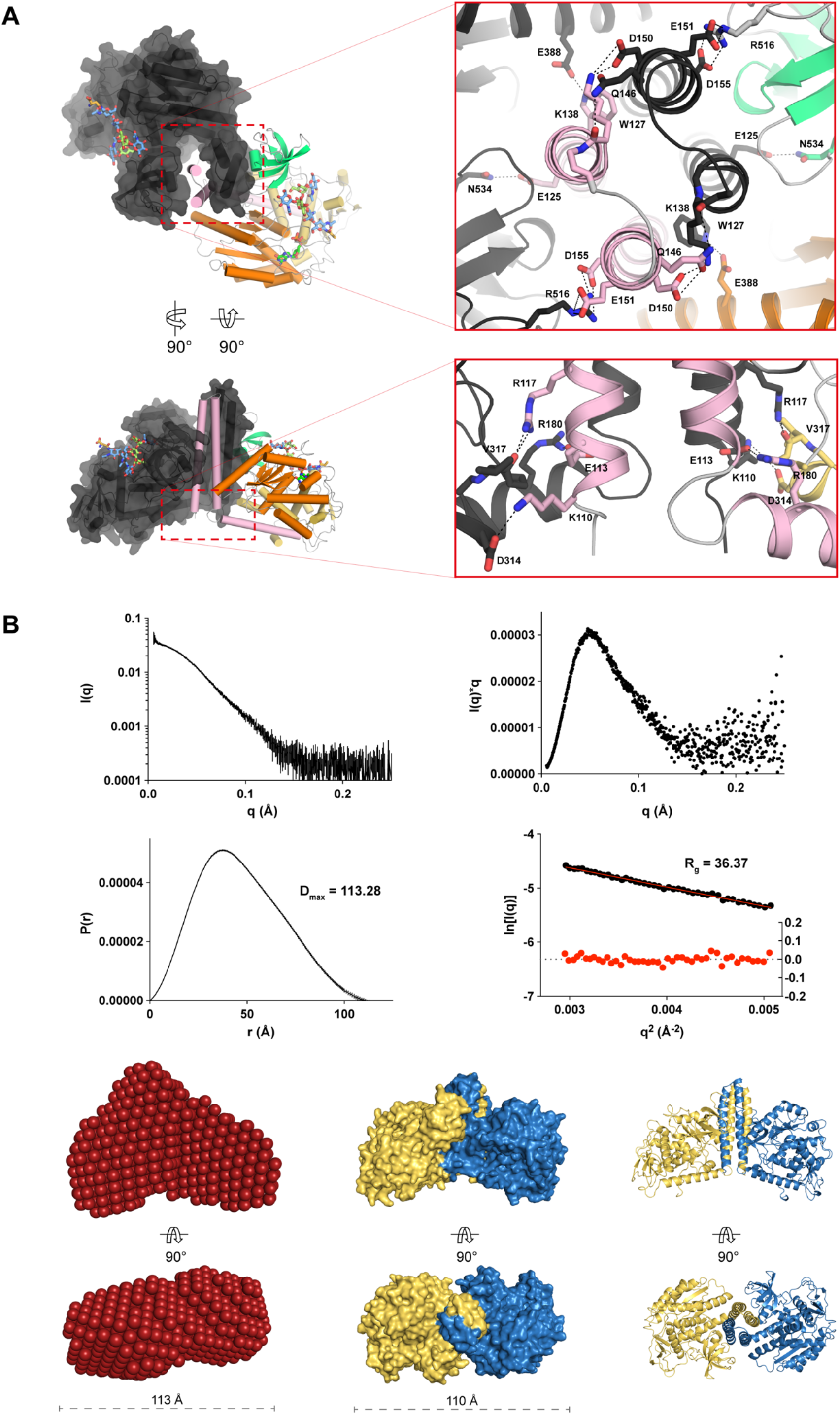
Interactions between the N-terminal coiled coil domains drive FUT8 dimerisation in solution. (A) Self-association of the N-terminal helices of FUT8 creates a four helix bundle that buries hydrophobic residues and creates multiple inter-chain salt bridges. (B) Intensity plot of FUT8 SAXS scattering (top left), Kratky plot derived from FUT8 scattering showing that it forms a compact particle in solution (top right), P(r) and Guinier plots indicating that FUT8 has a maximum dimension of 113.28 Å in solution (bottom left) and a radius of gyration of 36.37 Å (bottom right). (C) A bead model of FUT8 in solution generated from solution scattering data (left), corresponds well to the dimer observed in the FUT8 crystal structure shown as a surface (middle) and cartoon (right) view.

As mentioned, previous docking and molecular dynamics simulations were conducted based on the assumption that FUT8 is a monomer and suggested that the SH3 domain moves significantly after 20 ns of simulation^22^. Since the SH3 domain plays an important role in recognising the acceptor substrate, this would appear to be deleterious for catalysis. However, when considered as a dimer complex, the SH3-domain clearly binds the neighbouring chain’s N-terminal coiled-coil domain, which would appear to lock the SH3 domain in place (**Figure 4A**). To address this discrepancy, we performed MD simulations of unliganded FUT8 in both the monomeric and dimeric state over a period of 40 and 30 ns, respectively (**Figure 5**). Movements in the N-terminal coiled-coil domain and SH3 domain in the monomeric structure were replicated, with this conformational change enabling the burial of hydrophobic residues and disrupting the acceptor-binding surface of the enzyme. However, in the dimeric structure, no such movements were observed (Figure 5). In fact, for the dimer, the only significant motion observed on this time scale were in the active site loops A and B that encapsulate GDP. This data suggests that dimerisation of FUT8 is an important adaptation for buttressing the SH3 domains to maintain an extended bifurcated surface that can accomodate the bisected N-glycan acceptor substrate.

**Figure 5.**
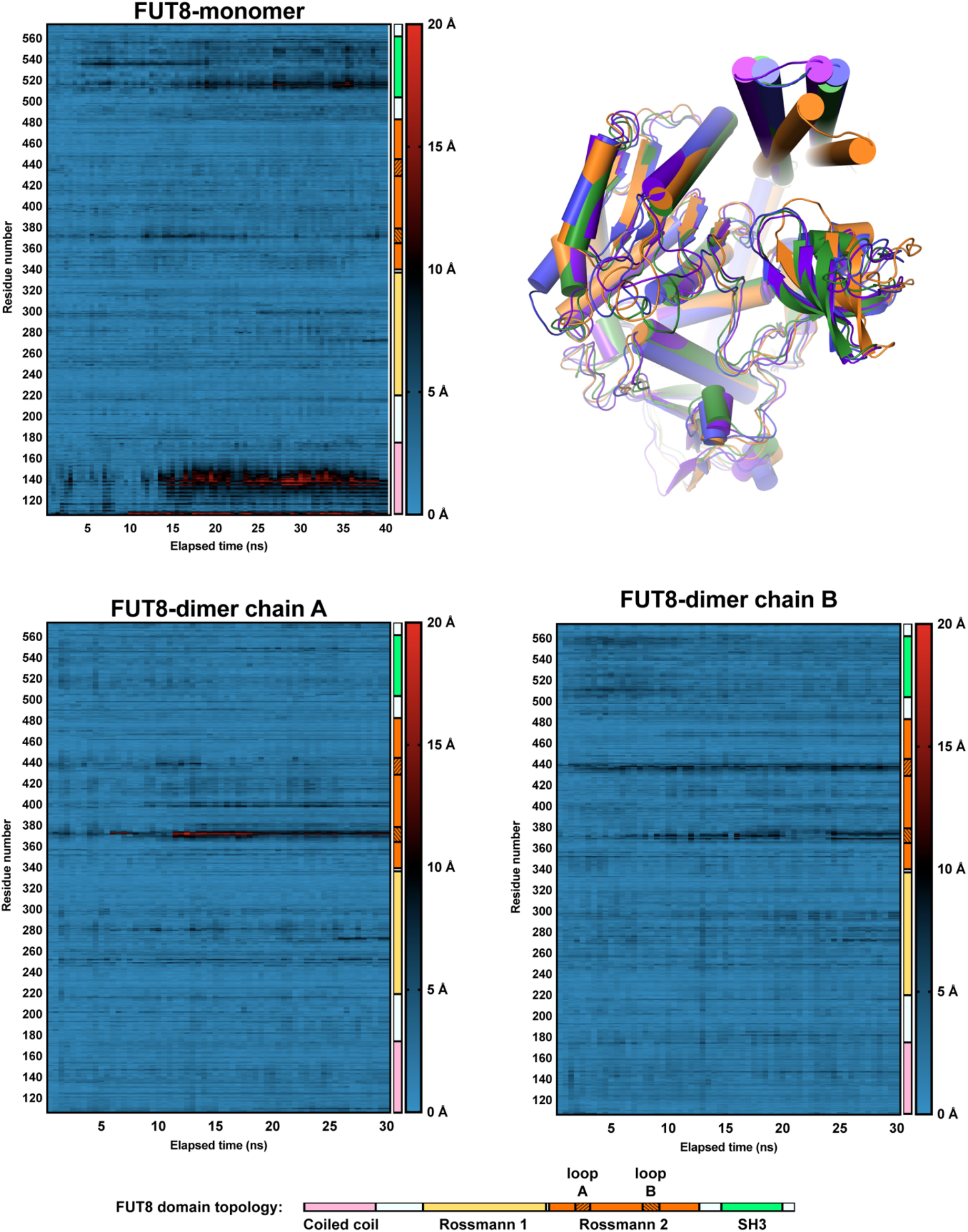
Heatmaps illustrating the displacement experienced by each amino acid in FUT8 during 30-40 ns of molecular dynamics simulation. The domain that each residue belongs to is illustrated for reference, with the shaded regions of the second Rossmann domain denoting the mobile GDP-binding loops A and B of the active site. An overlay of each monomer at t = 0 ns and the end points is provided in the top right (green is at t = 0 ns, orange is monomer at t = 40 ns, blue and purple are both dimer chains at t = 30 ns).

## Discussion

This collection of structures has revealed a unique conformational change in FUT8 associated with capturing GDP, and presumably its donor substrate GDP-Fuc. Arg365 plays a pivotal role in this process by forming a salt bridge with the beta-phosphate of GDP and Asp368/Arg441 of the mobile loops. The critical importance of this residue for catalysis is supported by previous mutagenesis studies^25^. Despite this unusual feature, once GDP is bound, its spatial orientation and the interactions it makes with FUT8 are largely the same as for other FUTs^26–30^. This observation is particularly relevant to those seeking to develop competitive inhibitors of FUTs: selectivity may be difficult to obtain with GDP mimics or molecular scaffolds that only interact with the GDP-binding site. On the other hand, these commonalities suggest that small molecule scaffolds may exist with pan-FUT inhibitory activity and that these might be adapted into selective inhibitors by exploring the acceptor binding site.

Original investigations into FUT8 mechanism by Ihara and co-workers indicated that FUT8 utilises a rapid equilibrium random mechanism^24^. This model postulates that substrates can bind independently to the enzyme in any order to form the Michaelis complex. The fact that we were able to obtain structures of FUT8 bound to acceptor or GDP alone, as well as the ternary complex, supports this mechanistic model. Our structures also reveal that there are no significant structural rearrangements associated with N-glycan binding and that the ordering of loops A and B upon GDP binding occur independently of the N-glycan binding site. Perhaps the greatest insights obtained from our structures, with respect to enzyme mechanism, is the realisation that the Glu273/Lys369 residue pair are the key catalytic residues. Our ternary complex clearly illustrates that Glu273 forms a close contact with the 6-hydroxyl group of the innermost GlcNAc of the N-glycan acceptor substrate, and that its basicity is modulated through interactions with Lys369. As such, Glu273 is the clear catalytic base residue. Concomitantly, Lys369 is able to shuttle a proton from Glu273 to the beta-phosphate of GDP, with which it forms a salt bridge, to facilitate departure of GDP from GDP-Fuc, enabling electrophilic migration of fucose onto the hydroxyl group nucleophile of the acceptor substrate.

With core-fucosylation playing such an important role in the function of proteins and the maturation of N-glycans, a great deal of effort has been invested in profiling the acceptor substrate preferences of this enzyme. Our structures provide a basis for understanding the vagaries of FUT8 substrate preference^31, 32^. What is clear from this work is that the SH3 domain of FUT8 plays a defining role in recognising N-glycan acceptor substrates. Modifications to the α3 branch of an N-glycan or a bisecting GlcNAc introduces steric clashes with the SH3 domain that prevents them from binding to FUT8. His535 of the SH3 domain also appears to form a crucial hydrogen-bond with the non-reducing end GlcNAc of N-glycan substrates. The importance of this SH3 domain to forming the acceptor substrate binding site necessitates its rigidity. Our molecular dynamics simulations support the hypothesis this requirement for rigidity is met by FUT8 dimerisation, there the N-terminal coiled-coil domains of one chain buttress the C-terminal SH3 domain of the other chain, to provide FUT8 with a rigid acceptor substrate binding site.

## Conclusion

FUT8 possess a unique method of capturing its GDP-Fuc donor substrate by using two mobile loops to encapsulate the nucleotide portion of this molecule: this process is largely driven by Arg365, which drives salt bridge formation between GDP and the two mobiles loops. This unique feature aside, FUT8 recognises GDP in much the same way as other FUTs, suggesting that all of these enzymes might be targetable with a common chemical scaffold. A ternary complex of FUT8, GDP, and N-glycan acceptor substrate revealed that Glu273 and Lys369 play a direct role in catalysis, with Glu273 acting as catalytic base, and Lys369 relaying a proton from Glu273 to the departing phosphate of the GDP-Fuc substrate. This complex also revealed the importance of the SH3 domain in providing a bifurcated surface for N-glycan recognition and in defining the acceptor substrate scope of FUT8. The importance of the SH3 domain in substrate binding appears to have driven the evolution of FUT8 as a dimer, which restricts the movement of the SH3 domain and stabilises the acceptor-binding subsite.

## Materials and Methods

### Cloning, expression, and purification of human and mouse FUT8

A gene encoding an N-terminal gp67 signal peptide, residues 105-585 of human FUT8 (UniProt ID: Q9BYC5) and a C-terminal His_10_ tag (**Table S1**) was synthesised and cloned into the pFastBac-1 vector (ThermoFisher) using *RsrII*/*XhoI*. A gene encoding an N-terminal gp67 signal peptide, V5 epitope tag, His_10_ tag, factor Xa site, and residues 68-585 of mouse FUT8 (UniProt ID: Q9WTS2) (**Table S1**) was synthesised and cloned into the pFastBac-1 vector (ThermoFisher) using *RsrII*/*XhoI*. Both constructs were expressed in Sf21 insect cells (Thermo Fisher) using the Bac-to-Bac Bacoulovirus Expression System (Thermo Fisher) by following the manufacturer’s protocol. Briefly, each plasmid was transformed into chemically competent DH10Bac *E. coli* cells (Thermo Fisher) and positive clones identified through a blue-white screen. Bacmid was prepared from these cells and transfected into Sf21 insect cells conditioned in Insect-XPRESS Protein-free Insect Cell Medium with L-glutamine (Lonza Ltd.) using the Cellfectin II reagent (ThermoFisher). Virus was passaged on Sf21 insect cells three times. For protein expression, 1 litre of Sf21 cells at a density of 1-2×10^6^ cells.ml^−1^ was infected with 30 ml of the P3 baculovirus and cultured at 27 °C for 72 h. The cells were pelleted by centrifugation (8,000×g, 20 min, 4 °C) and the supernatant collected. 10× buffer solution (112 ml, 500 mM Tris pH 7.5, 3M NaCl) was added to the supernatant before it was filtered through a 0.22 μm membrane. The buffered and filtered supernatant was passed through a pre-equilibrated 5 mL HisTrap Excel column (GE Healthcare). The column was washed with 20 column volumes of 50 mM Tris pH 7.5, 300 mM NaCl, followed by a one-step elution in 50 mM Tris pH 7.5, 300 mM NaCl, 500 mM imidazole to elute FUT8 from the column. Fractions containing protein, as judged by SDS-PAGE, were pooled and further purified by size exclusion chromatography (SEC) using a HiLoad 16/600 Superdex 200 column (GE Healthcare) equilibrated in 50 mM Tris pH 7.5, 150 mM NaCl. For MmFUT8_68-575_, the N-terminal tags were removed using Factor Xa protease (New England BioLabs) by incubating in 50 mM Tris pH 6.5, 150 mM NaCl, 2 mM CaCl_2_ overnight at room temperature. For HsFUT8_105-575_, the C-terminal His_10_ affinity tag was removed using Carboxypeptidase A (Merck) by incubating in 50 mM Tris pH 7.5, 150 mM NaCl overnight at room temperature. Protease-treated MmFUT8_68-575_ and HsFUT8_105-575_ were purified by running the reactions through a pre-equilibrated 1 mL HisTrap Excel column (GE Healthcare) and performing SEC on the column flow-through using a Superdex 200 Increase 10/300 GL column (GE Healthcare) equilibrated in 50 mM Tris pH 7.5, 150 mM NaCl. Concentration of the proteins was accomplished using Amicon centrifugal filters, NMWL 10 kDa (Merk-Millipore). Protein yields varied between batches but were always in the region of 5-10 mg per litre of cell culture.

### GDP-Glo assay of FUT8 activity

FUT8 activity was assayed using the GDP-Glo™ Glycosyltransferase Assay (Promega) with 3 μl reactions being conducted in a 1536-well microtiter plate. Reactions contained assay buffer (50 mM Tris pH 7, 100 mM NaCl, 0.01% Triton X-100 and 0.1% BSA), 5 μM A2SGP (Fushimi Pharmaceutical Co.), 10 μM GDP-Fuc and 5 nM FUT8, unless otherwise stated. After incubation for 20 min at room temperature, the reactions were stopped by the addition of 1 μl of 4% acetic acid prepared in assay buffer for 10 min. The resulting decrease in pH completely inactivated FUT8. To bring the pH back to neutral, 1 μl of 700 mM NaOH prepared in assay buffer was added to the reaction for 2 min. To detect the GDP product, 2.5 μl of GDP-Glo™ Glycosyltransferase Assay (Promega) nucleotide detection reagent was added. Plates were sealed and incubated at room temperature for 60 min. Chemiluminescence was quantitated on an EnVision Multimode plate reader (Perkin Elmer). Read-out time was 0.1 s per well. To determine the K_M_ of the A2SGP acceptor substrate under these conditions, reactions were conducted with serial dilutions of A2SGP starting at 80 μM, with 15 μM GDP-Fuc and 5 nM FUT8. To determine the K_M_ of the GDP-Fuc donor substrate under these conditions, reactions were conducted with serial dilutions of GDP-Fuc starting at 100 μM, with 5 μM A2SGP and 5 nM FUT8. All data was fit to the appropriate model using Prism 8 (GraphPad).

### Crystallisation of apo-MmFUT8_68-575_

MmFUT8_68-575_ in 50 mM Tris pH 7.4, 300 mM NaCl was concentrated to 15 mg.ml^−1^. A single crystal was grown over two weeks in a sitting drop at room temperature by mixing 0.5 μl well solution containing 0.25 M (NH_4_)_2_SO_4_ and 10% PEG3350 with 0.5 μl MmFUT8:GDP solution. The crystal was cryo-protected by supplementing the mother liquor with 25% glycerol and was cryo-cooled using liquid nitrogen.

### Crystallisation of MmFUT8_68-575_ in complex with GDP

MmFUT8_68-575_ in 50 mM Tris pH 7.4, 300 mM NaCl, 1 mM GDP at 0.1 mg.ml^−1^ was incubated at 4 °C overnight prior to concentration to a final of 15 mg.ml^−1^ MmFUT8_68-575_. A single crystal was grown over a week in a sitting drop at room temperature by mixing 0.5 μl well solution containing 0.25 M (NH_4_)_2_SO_4_ and 10% PEG3350 with 0.5 μl MmFUT8:GDP solution. The crystal was cryo-protected by supplementing the mother liquor with 25% glycerol and was cryo-cooled using liquid nitrogen.

### Crystallisation of apo-HsFUT8_105-575_

HsFUT8_105-575_ in 50 mM Tris pH 7.4, 50 mM NaCl was concentrated to 2 mg.ml^−1^. Crystals were grown over 3 days at 20 °C by mixing 1 μl well solution containing 12% (w/v) PEG 20000, 2.5% (v/v) DMSO and 0.1 M HEPES, pH 7.5 with 1 μl protein solution. The crystal was cryo-protected by supplementing the mother liquor with 25% ethylene glycol (v/v) and was cryo-cooled using liquid nitrogen.

### Crystallisation of HsFUT8 in complex with A2SGP and GDP

HsFUT8_105-575_ in 50 mM Tris pH 7.4, 50 mM NaCl, at 3 mg.ml^−1^ was mixed with A2SGP and GDP (in water) to final concentrations of 2 mg.ml^−1^ HsFUT8_105-575_, 0.5 mM A2SGP and 2 mM GDP. The mixture was incubated on ice for 30 min prior to setting-up crystallisation experiments. A single crystal was grown over 10 weeks in sitting drops at 8 °C by mixing 1 μl well solution containing 2 M NH_4_SO_4_, 0.2 M NaCl, and 0.1 M sodium cacodylate, pH 6.5, with 1 μl FUT8:A2SGP:GDP solution. The crystal was cryo-protected by supplementing the mother liquor with 3 M NH_4_SO_4_ and was cryo-cooled using liquid nitrogen.

### Data collection and structure determination

Data was collected at the Australian Synchrotron (MX2 beamline) and processed using XDS^34^. All structures were solved by molecular replacement using PHASER^35^ using the apo-structure of human FUT8 as a search model (PDB ID: 2DE0)^21^. The final models were built in Coot^36^ and refined with Phenix^37^. Data collection and refinement statistics are summarized in Table 1. The coordinates have been deposited in the Protein Data Bank (accession codes: 6VLD, 6VLE, 6VLF, and 6VLG). Figures were prepared using Pymol.

### Small Angle X-ray Scattering and Modelling

Size Exclusion Chromatography-Small Angle X-ray Scattering (SEC-SAXS) was performed using Coflow apparatus at the Australian Synchrotron^38, 39^. Purified HsFUT8 was analysed at a pre-injection concentration of 100 μM. Chromatography for SEC-SAXS was performed at 22 °C, with a 5/150 Superdex S200 Increase column, at a flow rate of 0.4 ml.min^−1^ in 50 mM Tris pH 7.9, 100 mM NaCl, 5% glycerol and 0.2 % sodium azide. The inclusion of glycerol and azide was essential to prevent capillary fouling due to photo-oxidation of buffer components. Scattering data were collected for 1 s exposures over a *q* range of 0.01 to 0.51 Å^−1^. A buffer blank for each SEC-SAXS run was prepared by averaging 10-20 frames pre- or post-protein elution. Scattering curves from peaks corresponding to HsFUT8 were then buffer subtracted, scaled across the elution peak, and compared for inter-particle effects. Identical curves (5-10) from each elution were then averaged for analysis. Data were analysed using the ATSAS package, Scatter and SOMO solution modeler^40^.

### Molecular dynamics

The FUT8 monomeric and dimeric systems were created using either one or two copies of chain A from the HsFUT8 substrate bound structure (6VLD). Each system was solvated in an orthorombic box, expanding 12 Å in each direction from the protein chain(s) and neutralised with Na^+^ and Cl^−^ at 150 mM. These steps were carried out with the AutoPSF Builder in VMD 1.9.4^41^. Molecular dynamics was simulated using the CHARMM36 force field for proteins^42^, the TIP3P^43^ water model, and sodium and chloride ion parameters from Benoit Roux and Coworkers^44^. Both systems were minimised with NAMD version 2.13^45^, using a conjugate gradient for 10,000 steps. Next, the systems were annealed by heating from 60 K to 300 K at a rate of 10 K/12 ps. After annealing, both systems were allowed to equilibrate at 300 K for 1,000 ps. The annealing and equilibration phases were carried out in a constant pressure/temperature (NPT) ensemble using the Langevin piston barostat set to 1 atm, and with harmonic constraints on all non-hydrogen protein atoms. After this the harmonic restraints on the protein were removed and the monomeric system was simulated for 40 ns and the dimeric system for 30 ns. All simulations were performed using a time step of 2 fs and using periodic boundary conditions with the Particle Mesh Ewald (PME) method to determine the electrostatics of the system. The aligned backbone RMSDs of the trajectories were calculated with the RMSDVT Visualizer Tool in VMD and plotted in Prism 8 (GraphPad).

## Acknowledgements

We thank the beamline staff at the Australian Synchrotron for help with X-ray data collection, as well as Dr. Janet Newman and Dr. Bevan Marshall at the Commonwealth Scientific and Industrial Research Organisation (CSIRO) Collaborative Crystallisation Centre (C3) for assistance in protein crystallization. This research was undertaken in part using the MX2 beamline at the Australian Synchrotron, part of ANSTO, and made use of the Australian Cancer Research Foundation (ACRF) detector. We would like to acknowledge support from: The Walter and Eliza Hall Institute of Medical Research; National Health and Medical Research Council of Australia (NHMRC) project grant GNT1139549; the Australian Cancer Research Fund; and a Victorian State Government Operational Infrastructure support grant.

## Author contributions

M.A.J. and M.D. performed structural studies; M.A.J. performed MD simulations; J.P.L, R.M. and A.J. produced recombinant protein; R.W.G. performed SAXS experiments and data analysis; E.D.G.-B. conceived the project; M.A.J. and E.D.G.-B. co-wrote the manuscript.

