## Supplementary Information for "Structural basis of substrate recognition and catalysis by fucosyltransferase 8"

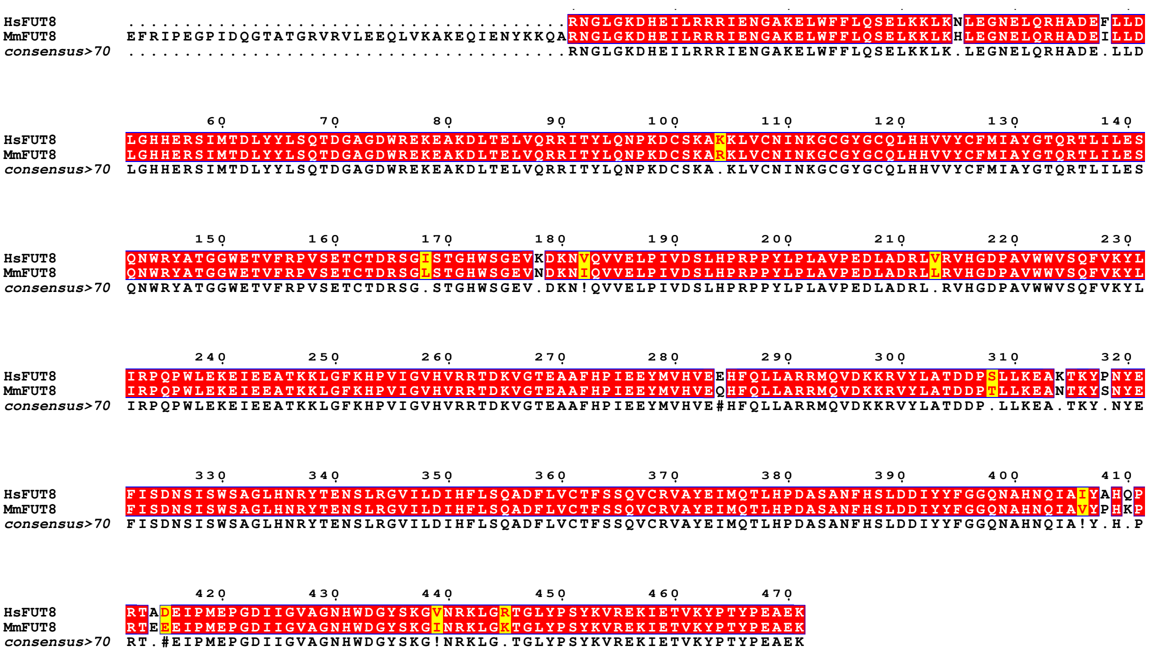


**Supplementary Figure 1.** Sequence alignment of $HsFUT8_{105\text{-}575}$ and $MmFut8_{68\text{-}575}$. The proteins are ≈97% identical over this region.

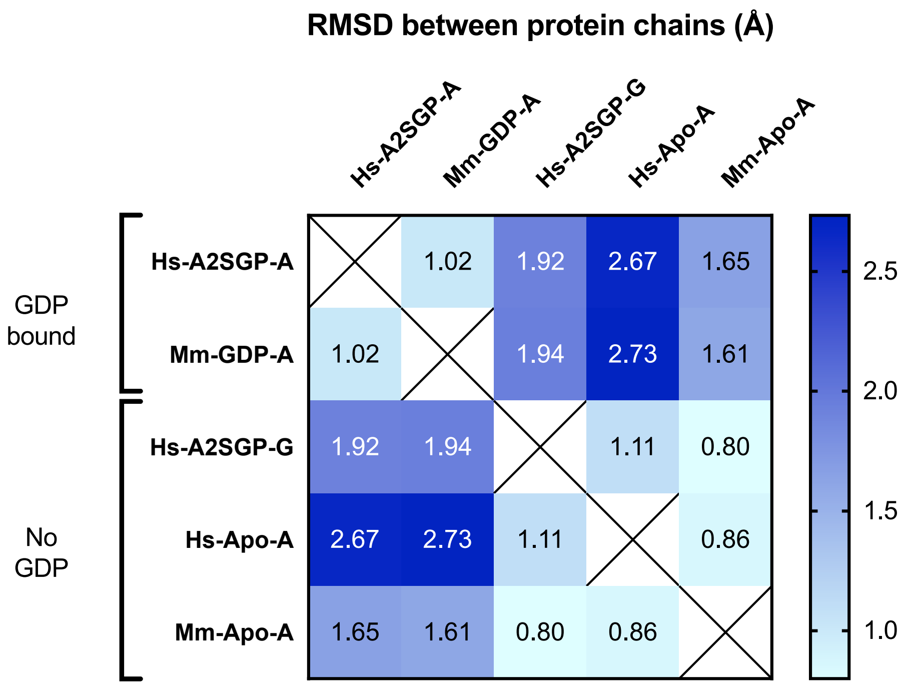


**Supplementary Figure 2.** Heatmap illustrating the RMSD between protein chains in the FUT8 structures reported in this study.

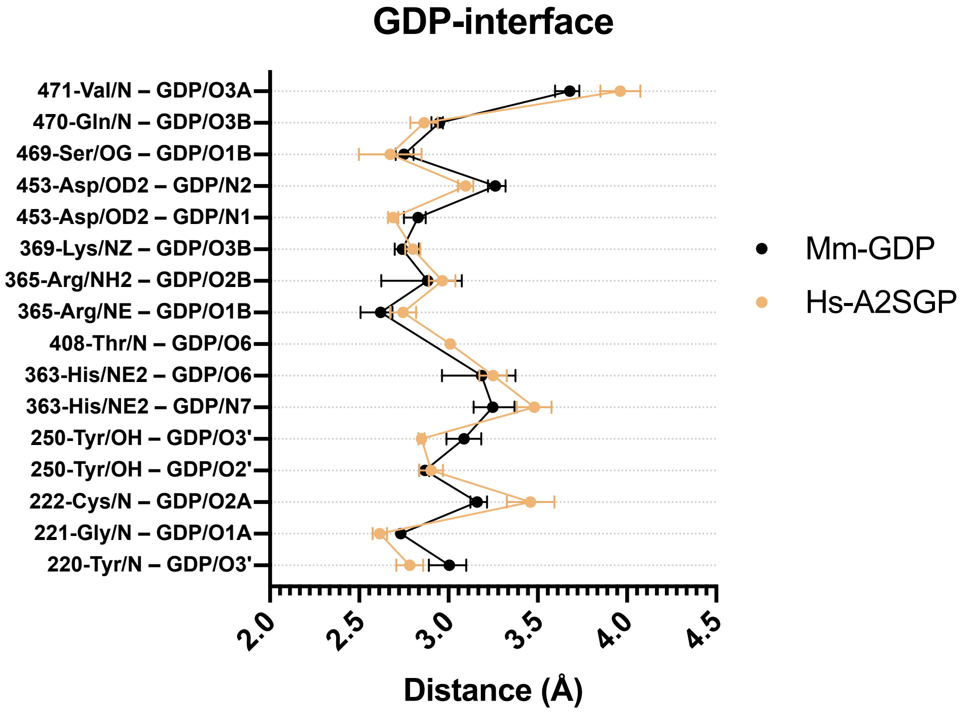


**Supplementary Figure 3.** A map of hydrogen bond distances between GDP and FUT8 for all relevant structures reported in this work.

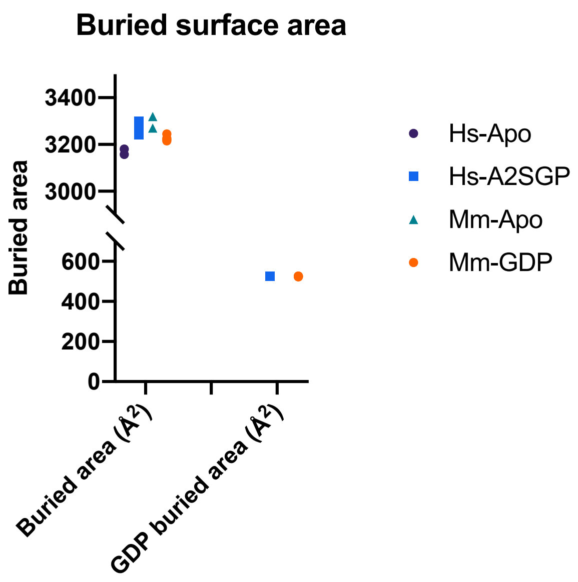


**Supplementary Figure 4.** Buried surface area of FUT8 upon dimerisation, and of GDP upon binding FUT8, for each of the structures reported in this work.

**Table S1.** The sequences of nucleotides and proteins used in this study. Restrictions sites used for cloning are underlined; signal peptides are shaded in grey; $His_{10}$ tags are shaded in yellow; V5 epitope tags are shaded in blue; and factor Xa sites are shaded in pink.

| protein | Synthetic dsDNA used for cloning | Protein sequence |
|---|---|---|
| HsFUT8 | GGATCTCGGTCCGAAACCATGCTACTAGTAAATCAGTCACACCAAGGCTTCA<br>ATAAGGAACACACAAGCAAGATGGTAAGCGCTATTGTTTTATATGTGCTTTT<br>GGCGGCGGCGGCGCATTCTGCCTTTGCGAGAAATGGTCTGGGGAAGGATCAT<br>GAAATCCTGAGGAGGAGGATTGAAAATGGAGCTAAAGAGCTCTGGTTTTTCC<br>TACAGAGTGAATTGAAGAAATTAAAGAACTTAGAAGGAAATGAACTCCAAAG<br>ACATGCAGATGAATTTCTTTTGGATTTAGGACATCATGAAAGGTCTATAATG<br>ACGGATCTATACTACCTCAGTCAGACAGATGGAGCAGGTGATTGGCGGGAAA<br>AAGAGGCCAAAGATCTGACAGAACTGGTTCAGCGGAGAATAACATATCTTCA<br>GAATCCCAAGGACTGCAGCAAAGCCAAAAAGCTGGTGTGTAATATCAACAAA<br>GGCTGTGGCTATGGCTGTCAGCTCCATCATGTGGTCTACTGCTTCATGATTG<br>CATATGGCACCCAGCGAACACTCATCTTGGAATCTCAGAATTGGCGCTATGC<br>TACTGGTGGATGGGAGACTGTATTTAGGCCTGTAAGTGAGACATGCACAGAC<br>AGATCTGGCATCTCCACTGGACACTGGTCAGGTGAAGTGAAGGACAAAAATG<br>TTCAAGTGGTCGAGCTTCCCATTGTAGACAGTCTTCATCCCCGTCCTCCATA<br>TTTACCCTTGGCTGTACCAGAAGACCTCGCAGATCGACTTGTACGAGTGCAT<br>GGTGACCCTGCAGTGTGGTGGGTGTCTCAGTTTGTCAAATACTTGATCCGCC<br>CACAGCCTTGGCTAGAAAAAGAAATAGAAGAAGCCACCAAGAAGCTTGGCTT<br>CAAACATCCAGTTATTGGAGTCCATGTCAGACGCACAGACAAAGTGGGAACA<br>GAAGCTGCCTTCCATCCCATTGAAGAGTACATGGTGCATGTTGAAGAACATT<br>TTCAGCTTCTTGCACGCAGAATGCAAGTGGACAAAAAAAGAGTGTATTTGGC<br>CACAGATGACCCTTCTTTATTAAAGGAGGCAAAAACAAAGTACCCCAATTAT<br>GAATTTATTAGTGATAACTCTATTTCCTGGTCAGCTGGACTGCACAATCGAT<br>ACACAGAAAATTCACTTCGTGGAGTGATCCTGGATATACATTTTCTCTCTCA<br>GGCAGACTTCCTAGTGTGTACTTTTTCATCCCAGGTCTGTCGAGTTGCTTAT<br>GAAATTATGCAAACACTACATCCTGATGCCTCTGCAAACTTCCATTCTTTAG<br>ATGACATCTACTATTTTGGGGGCCAGAATGCCCACAATCAAATTGCCATTTA<br>TGCTCACCAACCCCGAACTGCAGATGAAATTCCCATGGAACCTGGAGATATC<br>ATTGGTGTGGCTGGAAATCATTGGGATGGCTATTCTAAAGGTGTCAACAGGA<br>AATTGGGAAGGACGGGCCTATATCCCTCCTACAAAGTTCGAGAGAAGATAGA<br>AACGGTCAAGTACCCCACATATCCTGAGGCTGAGAAACACCACCATCACCAT<br>CACCATCACCATCACTGACTCGAGGCATG | MLLVNQSHQGFNKEHTSKMVSAIVL<br>YVLLAAAAHSAFARNGLGKDHEILR<br>RRIENGAKELWFFLQSELKKLKNLE<br>GNELQRHADEFLLDLGHHERSIMTD<br>LYYLSQTDGAGDWREKEAKDLTELV<br>QRRITYLQNPKDCSKAKKLVCNINK<br>GCGYGCQLHHVVYCFMIAYGTQRTL<br>ILESQNWRYATGGWETVFRPVSETC<br>TDRSGISTGHWSGEVKDKNVQVVEL<br>PIVDSLHPRPPYLPLAVPEDLADRL<br>VRVHGDPAVWWVSQFVKYLIRPQPW<br>LEKEIEEATKKLGFKHPVIGVHVRR<br>TDKVGTEAAFHPIEEYMVHVEEHFQ<br>LLARRMQVDKKRVYLATDDPSLLKE<br>AKTKYPNYEFISDNSISWSAGLHNR<br>YTENSLRGVILDIHFLSQADFLVCT<br>FSSQVCRVAYEIMQTLHPDASANFH<br>SLDDIYYFGGQNAHNQIAIYAHQPR<br>TADEIPMEPGDIIGVAGNHWDGYSK<br>GVNRKLGRTGLYPSYKVREKIETVK<br>YPTYPEAEKHHHHHHHHHH |
| MmFUT8 | GGATCTCGGTCCGAAACCATGCTACTAGTAAATCAGTCACACCAAGGCTTCA<br>ATAAGGAACACACAAGCAAGATGGTAAGCGCTATTGTTTTATATGTGCTTTT<br>GGCGGCGGCGGCGCATTCTGCCTTTGCGGCGGATCTTGGATCCCACCATCAT<br>CACCACCATCACCACCATCACGGCAAACCAATTCCCAACCCTTTGCTGGGAC<br>TGGATTCCACTATAGACGGCCGTGAATTCCGAATACCAGAAGGCCCCATTGA<br>CCAGGGGACAGCTACAGGAAGAGTCCGTGTTTTAGAAGAACAGCTTGTTAAG<br>GCCAAAGAACAGATTGAAAATTACAAGAAACAAGCTAGAAATGGTCTGGGGA<br>AGGATCATGAAATCTTAAGAAGGAGGATTGAAAATGGAGCTAAAGAGCTCTG<br>GTTTTTCTACAAAGCGAACTGAAGAAATTAAAGCATTTAGAAGGAAATGAA<br>CTCCAAAGACATGCAGATGAAATTCTTTTGGATTTAGGACACCATGAAAGGT<br>CTATCATGACAGATCTATACTACCTCAGTCAAACAGATGGAGCAGGGGATTG<br>GCGTGAAAAAGAGGCCAAAGATCTGACAGAGCTGGTCCAGCGGAGAATAACA<br>TATCTCCAGAATCCTAAGGACTGCAGCAAAGCCAGGAAGCTGGTGTGTAACA<br>TCAATAAAGGCTGTGGCTATGGTTGTCAACTCCATCACGTGGTCTACTGTTT<br>CATGATTGCTTATGGCACCCAGCGAACACTCATCTTGGAATCTCAGAATTGG<br>CGCTATGCTACTGGTGGATGGGAGACTGTGTTTAGACCTGTAAGTGAGACAT<br>GTACAGACAGATCTGGCCTCTCCACTGGACACTGGTCAGGTGAAGTAAATGA<br>CAAAAACATTCAAGTGGTCGAGCTCCCCATTGTAGACAGCCTCCATCCTCGG<br>CCTCCTTACTTACCACTGGCTGTTCCAGAAGACCTTGCAGACCGACTCCTAA<br>GAGTCCATGGTGACCCTGCAGTGTGGTGGGTGTCCCAGTTTGTCAAATACTT<br>GATTCGTCCACAACCTTGGCTGGAAAAGGAAATAGAAGAAGCCACCAAGAAG<br>CTTGGCTTCAAACATCCAGTTATTGGAGTCCATGTCAGACGCACAGACAAAG<br>TGGGAACAGAAGCAGCCTTCCACCCCATCGAGGAGTACATGGTACACGTTGA<br>ACAACATTTTCAGCTTCTCGCACGCAGAATGCAAGTGGATAAAAAAAGAGTA<br>TATCTGGCTACTGATGATCCTACTTTGTTAAAGGAGGCAAACACAAAGTACT<br>CCAATTATGAATTTATTAGTGATAACTCTATTTCTTGGTCAGCTGGACTACA<br>CAATCGGTACACAGAAAATTCACTTCGGGGTGTGATCCTGGATATACACTTT<br>CTCTCACAGGCTGACTTTCTAGTGTGTACTTTTTCATCCCAGGTCTGTCGGG<br>TTGCTTATGAAATCATGCAAACCCTGCATCCTGATGCCTCTGCGAACTTCCA<br>TTCTTTGGATGACATCTACTATTTTGGAGGCCAAAATGCCCACAATCAGATT<br>GCTGTTTATCCTCACAAACCTCGAACTGAAGAGGAAATTCCAATGGAACCTG<br>GAGATATCATTGGTGTGGCTGGAAACCATTGGGATGGTTATTCTAAAGGTAT<br>CAACAGAAAACTTGGAAAAACAGGCTTATATCCCTCCTACAAAGTCCGAGAG<br>AAGATAGAAACAGTCAAGTATCCCACATATCCTGAAGC**TGA**AAAATAGCTCG<br>AGGCATG | MLLVNQSHQGFNKEHTSKMVSAIVL<br>YVLLAAAAHSAFAADLGSHHHHHHH<br>HHHGKPIPNPLLGLDSTIDGREFRI<br>PEGPIDQGTATGRVRVLEEQLVKAK<br>EQIENYKKQARNGLGKDHEILRRRI<br>ENGAKELWFFLQSELKKLKHLEGNE<br>LQRHADEILLDLGHHERSIMTDLYY<br>LSQTDGAGDWREKEAKDLTELVQRR<br>ITYLQNPKDCSKARKLVCNINKGCG<br>YGCQLHHVVYCFMIAYGTQRTLILE<br>SQNWRYATGGWETVFRPVSETCTDR<br>SGLSTGHWSGEVNDKNIQVVELPIV<br>DSLHPRPPYLPLAVPEDLADRLLRV<br>HGDPAVWWVSQFVKYLIRPQPWLEK<br>EIEEATKKLGFKHPVIGVHVRRTDK<br>VGTEAAFHPIEEYMVHVEQHFQLLA<br>RRMQVDKKRVYLATDDPTLLKEANT<br>KYSNYEFISDNSISWSAGLHNRYTE<br>NSLRGVILDIHFLSQADFLVCTFSS<br>QVCRVAYEIMQTLHPDASANFHSLD<br>DIYYFGGQNAHNQIAVYPHKPRTEE<br>EIPMEPGDIIGVAGNHWDGYSKGIN<br>RKLGKTGLYPSYKVREKIETVKYPT<br>YPEAEK |
